# The influence of Schizotypy on Event-Related Oscillations in Sensory Gating during early Infant Development

**DOI:** 10.1101/2020.06.10.144014

**Authors:** Eleanor S. Smith, Trevor J. Crawford, Vincent M. Reid

## Abstract

Maternal schizotypic personality is thought to influence childhood risk for mental health and is a personality dimension elevated among schizophrenia-spectrum patients and their first-degree relatives, in whom neuro-oscillatory deficits have been observed. The current study investigated whether 6-month-old infants (*n* =46), and a subset of their biological mothers (*n* =34), who identified as either schizotypic (*n* =14) non-schizotypic (*n* =14), or an intermediate group (*n* =6), displayed reduced evoked-oscillatory activity. All mothers completed the Oxford-Liverpool Inventory of Feelings and Experiences as an index of schizotypy dimensionality. An auditory paired-tone paradigm was used to probe oscillatory activity, revealing that although the infants’ evoked-oscillations displayed differences between *Stimulus 1* and *2*, there were no group differences between infants of schizotypic and control mothers. Their mothers, however, displayed differences, with reduced amplitudes toward *Stimulus 1* in schizotypic mothers; consistent with literature on early sensory processes, showing sensory gating is impaired in schizophrenia-spectrum disorders.

## Introduction

Maternal personality is known to play an influential role in determining childhood risk factors for mental health. For example, specific maternal psychopathologies have been associated with P50 sensory gating abilities, illustrated through event-related potentials (Ross and Freedman, 2015). Atypical P50 sensory gating is an established biological trait of schizophrenia (Raine, 2006), which is also observed in individuals with schizotypal personality disorder (Cadenhead et al., 2000) and schizotypic personality traits among those who remain in the general population (Smith et al., 2019). These deficits are also observed in infants and children of parents with psychoses or severe anxiety disorders (Ross and Freedman, 2015) as well as those at risk of developing schizophrenia-spectrum disorders (Cadenhead, Light, Shafer, and Braff, 2005). The presence of this deficit amongst all capacities of the schizophrenia-continuum supports sensory gating as a potential biomarker for the general risk of psychopathology.

The sensory gating endophenotype is described as having the potential to extend into infancy (Freedman et al., 2002) and to shape the development of personality into adulthood (Corr, 2010). The core neuropsychological and neuro-oscillatory dysfunctions associated with sensory gating abnormalities, however, are unresolved despite their predictive capabilities of identifying potential future psychopathology (Debbané and Barrantes-Vidal, 2015). In the present research, a 6-month-old infant population was chosen due to the developmental trajectories observed in the existing sensory gating literature (for example, Smith et al., 2019). We know from the literature that intact sensory gating can be observed during periods of REM sleep from 2-(Hutchison et al., 2017) or 3-months of age (Hunter et al., 2015), although there are inconsistencies in developmental trajectories due to large age-gaps amongst published literature (for example, Hunter et al., 2015: 3-months and then 4-years). Kisley et al. (2013) explored ‘typical’ sensory gating abilities among infants between the ages of 1-5-months; illustrating that over this period, sensory gating suppression ratios and spectral power both significantly correlated with age (*r* = −0.77, *n* = 11, *p* < 0.01, *r* =0.70, *n* = 11, *p* < 0.05, respectively). We could therefore confidently explore 6-month-old infants with the expectation of observing their sensory gating ability during periods of sleep. The infant portion of the present paradigm was therefore carried out while the infants were sleeping. The rationale for this is based on research by Hunter et al. (2015), who explored sensory gating among sleeping children (~4-years-old). Previous research had verified that like adults, REM sleep was a favourable state in which to measure P50 sensory gating in infants as movement artefacts are reduced (Kisley et al., 2003).

As far as we are aware, this is the first time the influence of maternal disposition/personality on sensory gating ability during infancy is being explored in electrophysiological oscillatory activity. Despite this novelty, Huggenberger et al. (2013) explored a different inhibitory mechanism, pre-pulse inhibition, among 4-day-old and 4-month-old infants in connection with their mothers’ disposition. It was concluded that infants of mothers reporting less ‘social recognition’ and higher ‘social isolation’ developed less pre-pulse inhibition during the four months after birth than did infants of mothers with higher ‘social recognition’ and less ‘social isolation’. This confirms that parental disposition impacts inhibitory abilities of their infants at 4-months. With the knowledge that parental disposition can influence infants inhibitory abilities at 4-months, it is of fundamental interest to explore the different frequencies observed across the electrophysiological spectrum (for example beta and gamma), associated with auditory sensory gating within the non-clinical realm of the schizophrenia-spectrum and to explore whether these endophenotypic deficits are observed earlier in the developmental trajectory.

The inhibition of responses to irrelevant stimuli is an essential cognitive ability for humans in everyday life. The ability to ‘gate out’ irrelevant stimuli is known as *sensory gating*, which can be demonstrated as an attenuated neural response to the second identical stimulus in a paired-tone paradigm and is considered an automatic inhibitory function (Freedman et al., 1987). In the auditory modality, the paired-tone paradigm: two tones presented in close succession, have been widely applied across the schizophrenia-spectrum (Patterson et al., 2008). The P50 auditory event-related potential (ERP) response to the second stimulus (*S2)* is typically reduced compared with that of the first stimulus (*S1*). This suppression, termed P50 gating, is thought to serve as a protective mechanism against flooding of the higher-order cortical centres with unnecessary information (Braff et al., 2007), with schizophrenia patients and a proportion of their clinically unaffected relatives exhibiting reduced P50 ERP suppression.

Most studies of auditory sensory gating rely on information obtained by amplitude measurements of specific peaks in the time domain, for example the P50 event-related potential component (ERP; for example, Park et al., 2015; Smith et al., 2019); however, the auditory stimulus used to elicit P50 ERPs also evokes brain activity in a wide spectrum of frequency bands. There is evidence that at least two broadly defined frequency bands, each associated with a distinct role in information processing, contribute to these auditory evoked response amplitudes (Clementz and Blumenfeld, 2001). Auditory P50 ERPs overlap morphologically with the evoked gamma frequency (20-50 Hz; Clementz and Blumenfeld, 1997; Muller et al., 2001) which is associated with the initial neural registration of a sensory stimulus (Karakas and Basar, 1998). Although the P50 ERP is thought to be a subcomponent of gamma frequency activity (Basar et al., 1987; Clementz et al., 1997), auditory suppression deficits in the schizophrenia-spectrum might, to an even greater extent, reflect poor activation of a low frequency response occupying the 1-30Hz range (Clementz and Blumenfeld, 2001). This range spans the beta band, which is associated with the detection of salient changes in sensory stimuli (Haenschel et al., 2000) and is thought to signify attentional engagement to task-relevant features of stimulus processing. Response to *S1* in the low beta frequency range (13-30 Hz), which is involved in the attention/encoding process and has been negatively associated with P50 gating to *S2* stimuli (Hong et al., 2004a). These studies suggest that gamma-band (30-50 Hz) and beta-band oscillations (13–30 Hz) contribute to auditory P50 ERP responses (see Uhlhaas and Singer, 2010, for a review), although the precise mechanisms remain to be determined.

Paired-tone suppression appears deficient in the schizophrenia-spectrum in the form of smaller evoked responses to *S1* in fronto-central regions (Freedman et al., 1987; Johannesen et al., 2005), rather than larger, less effectively “gated” responses to *S2* (Johannesen et al., 2005). For example, Johannesen et al. (2005) found that less suppression was observed by schizophrenia patients in the low frequency response, and that the response to *S1* was reliably smaller with no evidence of higher, poorly gated, responses to *S2*. Therefore, in conjunction with results of other research demonstrating lower *S1* but normal *S2* amplitudes in schizophrenia at P50 (e.g., Boutros et al., 1991, Patterson et al., 2000, Zouridakis et al., 1997) and low frequency responses (Blumenfeld and Clementz, 2001), these results cannot be fully explained by the predominant recurrent inhibition model of sensory gating. In addition, significant correlations were observed between gamma and beta power to both *S1* and *S2* stimuli, between P50 amplitudes and beta power to both *S1* and *S2* stimuli, and between P50 *S2* amplitude and gamma *S1* power (Hall et al., 2011). These results replicate those of Kisley and Cornwell (2006) who reported positive correlations between P50 amplitudes and beta/gamma power to *S1* and *S2* stimuli, and those of Hong et al. (2004b) who documented positive correlations between gamma and beta responses. The positive relationship between beta and P50 responses to both *S1* and *S2* stimuli (correlations between .34 and .45) suggest a contribution of beta to P50 ERP activity in general and the positive relationship between *S1* gamma and *S2* P50 amplitudes (*r* = .45) suggest a contribution of gamma *S1* activity to P50 *S2* ERP too. These results are consistent with the notion that both gamma and beta EROs contribute to the generation of P50 ERPs.

It is important to decompose the specific frequency components of ERPs because each event-related oscillation (ERO) is thought to reflect different aspects of early sensory information processing (Kopell et al., 2000; Traub et al., 1999; Traub et al., 1999; Basar-Eroglu et al., 1996). For example, beta activity has been implicated in novelty detection and salience encoding, providing recognition that a novel or salient stimulus warrants monitoring (Uhlhaas et al., 2008; Kopell et al., 2000; Kisley and Cornwell, 2006), whereas gamma oscillations are associated with the immediate registration of sensory stimuli (Kopell et al., 2000; Tallon-Baudry and Bertrand, 1999) reflecting the “binding of spatially distinct neuronal activities within and between brain regions” necessary for coherent sensory perception and processing (Uhlhaas et al., 2008; Singer and Gray, 1995). Deficits in both frequency ranges would result in weak neuronal signals and propagation across broadly distributed networks, potentially leading to impaired perceptual experiences and cognitive dysfunction (Kopell et al., 2000; Traub et al., 1999; Basar-Eroglu et al., 1996; Tallon-Baudry and Bertrand, 1999).

The literature is mixed as to whether similar deficits in oscillatory responses to P50 are observed in first-degree relatives. For example, Hong et al. (2008) suggested that the first-degree relatives of those diagnosed with schizophrenia displayed no difference in oscillatory gating compared to controls (Hong et al., 2008). Impairments in sensory gating are thought to be predictive of parent-reported problems with attention, anxiety, and externalizing problems at preschool age (Hutchison et al., 2013). Identifying infant brain changes associated with the later onset of ‘symptoms’ or traits homogenous to schizotypy (or other sub-clinical manifestations of the schizophrenia-spectrum) suggest that, at least in part, these may be due to alterations in brain development that occur years prior to clinical, or even sub-clinical presentation. Smith et al. (2019) identified correlational relationships illustrating that having a mother with high-schizotypic traits increased their 6-month-old infants’ suppression ratios, implying a similar sensory gating deficit to that observed in their mothers. Identification of early brain changes may therefore be useful in identifying infants who would benefit from primary prevention interventions and would assist in the uncovering of earlier aberrant brain processes where behaviours were not yet observable.

A number of studies have taken a psychometric approach to further understand and delineate the relationship between schizotypy and P50 suppression (Hall et al., 2011). In general, research has found that high levels of schizotypy were associated with poor P50 suppression (Croft et al., 2004; Wan et al., 2006; Smith et al., 2019), supporting the idea that schizotypic individuals display similar deficits to those found in patients with schizophrenia and individuals at high-risk for psychosis (Ettinger et al., 2014; Cohen et al., 2015). Although, the research differs in terms of which dimensions of schizotypy are associated with this deficit. Croft et al. (2001) found the unreality dimension of schizotypy was associated with atypical sensory gating patterns, but other literature has suggested it to be the negative dimension (Wang et al., 2004), positive dimension (Croft et al., 2001, 2004), or the cognitive disorganisation dimension rather than distinctly positive or negative symptoms of schizotypy (Evans et al., 2007; Park et al. 2015). Moreover, reduced P50 suppression in participants high in cognitive disorganisation is consistent with the correlation of other measures of gating, such as pre-pulse inhibition; a phenomenon referring to the reduction of the startle response if a startling stimulus is preceded by a weaker stimulus (Graham, 1975). Similarly, high scores of cognitive disorganisation have been associated with reduced pre-pulse inhibition (Evans et al., 2005). Thus, it appears that impairments in the ability to inhibit extraneous stimuli are associated with experiences of disorganisation (Evans et al., 2007).

The primary aim of the current research was to observe whether mothers who identify as exhibiting schizotypic traits demonstrate reduced evoked amplitudes in their neuro-oscillatory responses during a sensory gating paradigm (*Experiment 1*). As stated previously paired-tone suppression appears deficient in the schizophrenia-spectrum in the form of a smaller evoked response to S1, rather than larger, less effectively “suppressed” responses to S2, as seen in Smith et al. (2019). We therefore hypothesised that the mothers with schizotypy would illustrate reduced oscillatory amplitudes in the gamma and beta frequency ranges in response to *S1*. We also wanted to explore the dimensional nature of the sensory gating abnormality; predicting that those individuals exhibiting high schizotypy would display poorer sensory gating than those who experience low schizotypy scores. It was also hypothesised that poor sensory gating would be associated with higher manifestations of cognitive disorganisation and impulsive non-conformity, based on previous literature by Park et al. (2015) amongst other research.

The present study aims to explore whether these oscillatory sensory gating responses have similar manifestations in the 6-month-old offspring of the maternal cohort (*Experiment 2*). This question was asked to explore individual differences in sensory gating abilities, in this case whether abnormalities are observable at 6-months, or whether differences are manifested later in development. Current infant literature investigating the influence of schizotypy on sensory gating development is sparse; preventing the extent to which we can state that the early manifestation of schizotypic traits could be considered as a precursor to mental illness. Correlational relationships between maternal schizotypy and P50 gating have been identified (Smith et al., 2019) and maternal disposition has been shown to impact inhibition in their 4-month-old offspring (Huggenberger et al., 2013); suggesting parental personality influences such inhibitory abilities from early in development. For this reason, it is important we explore infant electrophysiological changes that have the potential to be associated with the later onset of traits.

## Materials and Methods

### Experiment 1: Adult Cohort

#### Participants

Thirty-five mothers (*M age* = 32.9 years, *SD* = 3.6 years; all white-Caucasian) participated in the research and were recruited from the Lancaster University Department of Infant and Child Development database. All participants were from a non-clinical population but completed the Oxford-Liverpool Inventory of Feelings and Experiences-Short Form (sO-LIFE; Mason, Linney, and Claridge, 2005), which is a four-scale questionnaire used to measure psychosis-proneness, principally schizotypy, in healthy individuals in the general population (Mason and Claridge, 2006). Thirty-four mothers were included in the final analysis; including fourteen participants who identified as a schizotypic mother (SZTm), fourteen controls (CONm) and six participants whose sO-LIFE total score did not fall into one of the two categories. These participants were excluded from the group analyses. One participant was excluded from the final sample due to poor data quality. All mothers who participated in *Experiment 1* (*n*=34) were a subset of the infants’ mothers from *Experiment 2* (*n*=75). See Table 3 for a breakdown of how many participants were included in each analysis and how many identified as SZTm or CONm.

The mean of the sO-LIFE total score across the population was calculated (total *M*=10.07, total *SD*=6.77) and the SZTm condition was determined by the *M*+.5*SD* (sO-LIFE Scores>13.46) and the CONm condition by the *M*-.5*SD* (sO-LIFE Scores 6.68>0.0). This criterion was used as a result of its previous use in the schizotypy literature (for example, Park et al., 2015). The mean sO-LIFE score among the SZTm was 15.60 (SD=2.82, Range = 12-21) and among the CONm the mean sO-LIFE score was 3.36 (SD=1.55, Range = 0-5). Ethical approval was obtained with the Lancaster University Ethics Board in accordance with the Declaration of Helsinki and the North West – Lancaster Research Ethics Committee for the NHS. All participants gave written informed consent for the study.

### Experiment 2: Infant Cohort

#### Participants

Seventy-five non-related infants, aged 6-months (*M*=5.9 months; *SD*=8.77 days; 39 female; all white-Caucasian) participated in the study, a subset of whom were the biological offspring of the previously discussed maternal cohort (n=34). To clarify, seventy-five infants participated in *Experiment 2*, of which, all their Mothers completed the sO-LIFE questionnaire (Mason, Linney, and Claridge, 2005), but only a subset of their mothers had participated in *Experiment 1*. A further twenty-eight were excluded from the final sample due to: poor data quality (*n*=14), incomplete sO-LIFE questionnaires (*n*=2), or the participant did not produce over 20% good epochs (*n*=13). A small group of participants’ sO-LIFE scores did not identify with either control or schizotypic groups (*n*=9), thus they were not included in the group analysis. Thirty-seven (20 male) infants were included in the final analysis (sO-LIFE Score total *M*=8.15, total *SD*=6.26). The final sample included fifteen participants identifying as an infant of a schizotypic mother (iSZTm; determined by the *M*+.5*SD* (sO-LIFE Scores>11.28)) and the remaining twenty-two participants were infants of control mothers (iCONm; determined by *M*-.5*SD* (sO-LIFE Scores 5.02>0.0)). The mean sO-LIFE score among the iSZTm was 15.93 (SD=3.88, Range = 12-28) and among the iCONm the mean sO-LIFE score was 3 (SD=1.51, Range = 0-5). All participants were from a non-clinical population. See Table 3 for a breakdown of how many participants were included in each analysis and how many identified as iSZTm or iCONm.

In respect to the present attrition rate, for an EEG experiment with infants, this is a typical sample size for studies using these methods (e.g., Stets et al., 2012; Smith et al., 2019) or substantially greater than the sample size for studies on schizotypy during development (Hunter et al., 2015). Ethical approval was obtained with the Lancaster University Ethics Board in accordance with the Declaration of Helsinki and the North West – Lancaster Research Ethics Committee for the NHS. All parents of infant participants gave written informed consent for the participation of their infant in the present study.

An independent samples t-test and Chi-squared analysis explored whether group differences were observed for the variables: age, gender, mothers mental health experiences, family history of mental health, or birth complications. The analyses presented no significant differences between schizotypic and control groups in both infant (Table 1) and maternal cohorts (Table 2). No significant differences were observed between the groups.

**Table 1.** Table indicating similarities between the iSZTm and iCONm groups in their demographic information, as collected using a general information questionnaire.

|  | Infants of Mothers<br>with Schizotypy<br>(*n*=15)<br>*M*(*SD*) | Infants of Control<br>Mothers<br>(*n*=22)<br>*M*(*SD*) | *p*-<br>value | Chi-<br>square |
| --- | --- | --- | --- | --- |
| Infant Age (days) | 179.27 (9.51) | 178.52 (7.77) | .793 |  |
| Infant Gender |  |  |  | .476 |
| Female | *n*= 6 | *n*= 11 |  |  |
| Male | *n*= 9 | *n*= 11 |  |  |
| Mothers Mental Health Experiences | 1.60 (.51) | 1.86 (.35) | .070 |  |
| Family History of Mental Health | 1.53 (.52) | 1.59 (.50) | .737 |  |
| Birth Complications | 2.00 (1.00) | 1.57 (.84) | .158 |  |

**Table 2.** Table indicating similarities between the SZTm and CONm groups in their demographic information, as collected using a general information questionnaire.

|  | Mothers with<br>Schizotypy<br>(*n*=14) | Control Mothers<br>(*n*=14) | *p*-value | Chi-<br>square |
| --- | --- | --- | --- | --- |
| Mothers Age (years) | 32.08 (4.89) | 32.50 (2.02) | .77 |  |
| Mental Health Experiences | 1.58 (.51) | 1.93 (.27) | .10 |  |
| No Mental Health Experience | *n*= 7 | *n*=13 |  | .10 |
| Depression | *n*= 1 | *n*= 0 |  |  |
| Anxiety | *n*= 1 | *n*= 1 |  |  |
| Postnatal Depression | *n*= 2 | *n*= 0 |  |  |
| Depression/Anxiety | *n*= 3 | *n*= 0 |  |  |
| Family History of Mental Health | 1.50 (.52) | 1.71 (.47) | .28 |  |

**Table 3.** Table outlining the number of participants whose data was included in each element of the analysis.

|  | *n* | Schizotypic : Control :<br>No group |
| --- | --- | --- |
| ***Experiment 1: Maternal Cohort*** |  |  |
| Total Mothers Participated in Research | 35 |  |
| Mothers included in Repeated-Measures ANOVA Analysis | 28 | 14 : 14 |
| Mothers included in Pearson's Correlational Analysis | 34 | 14 : 14 : 6 |
| ***Experiment 2: Infant Cohort*** |  |  |
| Total Infants Participated in Research | 75 |  |
| Infants included in Repeated-Measures ANOVA Analysis | 37 | 15 : 22 |
| Infants included in Pearson's Correlational Analysis | 46 | 15 : 22 : 9 |

#### Procedure

Prior to participation, the caregiver completed a series of questionnaires, which included the Oxford and Liverpool Inventory of Feelings and Experiences (sO-LIFE; Mason, Linney, and Claridge, 2005) as a measure of schizotypy dimensionality. The EEG cap was soaked in a warm water, sodium chloride solution and baby shampoo before fitting to the participant’s head prior to the infant falling asleep or the mother being made comfortable in a darkened room. Once fitted and following confirmation that each electrode responded to electrical activity, the trial procedure began. The auditory stimuli was presented 80-centimetres away, between 70-77dB, with an average of 75dB in serial presentation until the 15-minutes were complete, the infant woke or became restless. The infant was then left to complete their natural sleep period. Throughout the testing period the participant’s status was video-recorded to index activity. The biological mothers of the infant cohort were invited back to participate in the same paradigm at a later date in a darkened room in a resting state.

### Materials and Stimuli

The materials, stimuli, and analysis utilised were the same for *Experiment 1* and *Experiment 2* with the exception of the scalp regions and time windows for analysis. All participants experienced a pair of single-sound stimuli that were based on Park, Lim, Kirk, and Waldie (2015). See Figure 1 for a schematic of the paired-tone paradigm. Two identical tones of 50ms length (*Stimulus 1: S1*, and *Stimulus 2: S2*) were presented with a 500ms inter-tone interval every 10-seconds for 15-minutes or until the infant woke. The paired tones were presented between 70-77dB, with an average presentation of 75dB and had a tonal quality of 1000Hz.

**Figure 1.**
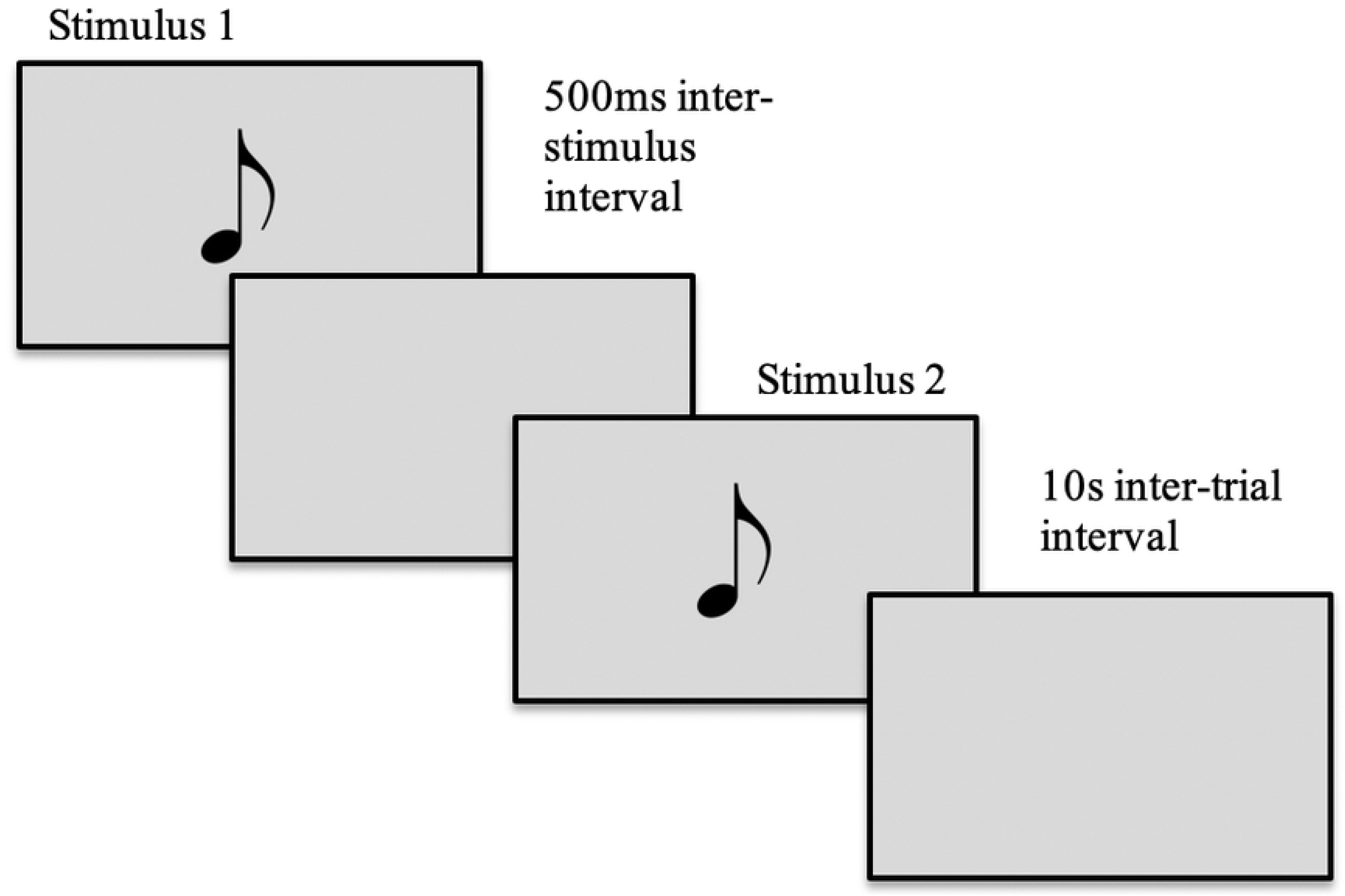
A graphical representation of the paired-tone paradigm. The tones were presented between 70-77dB with a tonal quality of 1000Hz.

The maternal cohort experienced a range of 56–64 paired-stimuli repetitions: equating to a minimum of 11–12 good trials, and contributed an average of 49.48 (88.2% artefact free trials; SD = 5.35; range = 42–61) artefact-free trials for *S1*, and on average 49.58 (88.3% artefact free trials; SD = 5.12; range = 41–62) artefact-free trials for *S2*. A paired-samples *t*-test displayed no significant differences in trial numbers between *S1* and *S2* (*t*(34)=−.21, *p*=.84). The infants experienced a range of 63-141 paired-stimuli repetitions, dependent on length of sleep, and contributed an average of 48.18 (51.62% artefact-free trials; SD= 24.74; Range: 23-127 for *S1*, and on average 48.20 (52.04% artefact-free trials; SD= 24.01; Range: 22-126) for *S2*. There were no significant differences in trial numbers between *S1* and *S2* (*t*(34)=.98, *p*=.33).

All electrophysiological signals were recorded using Electrical Geodesics Inc. amplifiers (input impedance=80KΩ; sampling rate=500 Hz) and event-related oscillations were measured using an EGI Hydrocel GSN-128 electrode 1.0 net and pre-processed using Netstation 4.5.4. All EEG signal processing and analyses were performed blind to group membership. The data was 0.5-65Hz bandpass filtered and segmented to create epochs from 400ms before to 1200ms after stimulus-onset for each trial. For the elimination of electrical artefacts caused by eye and body movements, data was rejected offline by the visual editing of trial-by-trial data. This was carried out in conjunction with an artefact detection toolset in Netstation, which highlighted whether a channel was ‘bad’ for more than 80% of the recording, which was determined by a threshold of 200μV to remove outlier values resulting from artefacts, or if they contained more than 12 bad channels in a trial. Participants required a minimum of 20% good trials for each stimulus to be included in further analyses.

Based on previous literature (Smith et al., 2010; Popov et al., 2011a) and visual inspection of the grand and individual means, we selected the scalp area, time window and frequency band for the maternal and infant cohorts. For the maternal cohort, gamma and beta frequencies were measured over the left-frontal (the average of channels 12, 18, 19, 20, 22, 23, 24, 26, 27), and right-frontal (the average of channels 3, 4, 5, 9, 10, 11, 118, 123, 124) regions, in the 50-250ms time window (Figure 2). Beta activity was analysed between 13-20Hz and gamma between 35-50Hz. For the infant cohort, gamma and beta frequencies were measured over the left-temporal (the average of channels 47, 51, 52), right-temporal (the average of channels 115, 116), and left-frontal (the average of channels 12, 20, 24) regions, in the 100-325ms time window (Figure 2). Beta activity was analysed between 10-20Hz and gamma between 30-50Hz.

**Figure 2.**
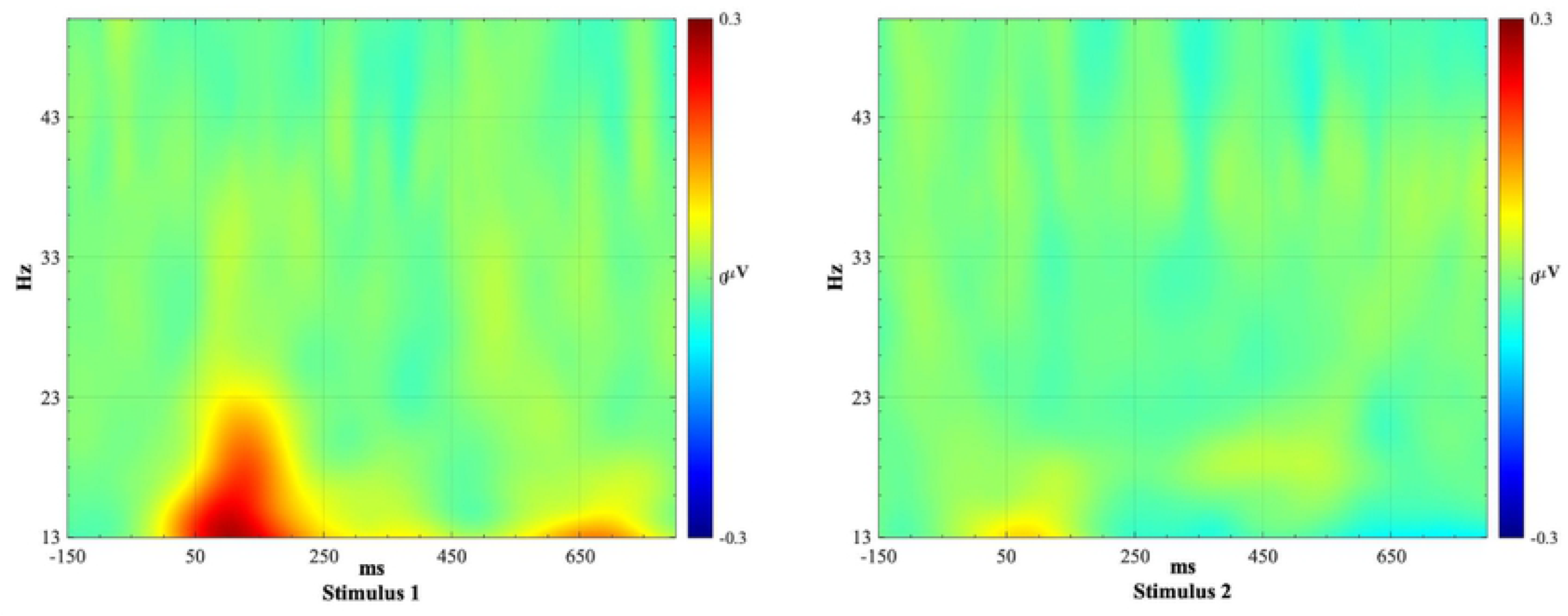
Maternal and Infant regions of interest, which were averaged across the 50-250ms and 100-325ms time windows, respectively.

The artefact-free segments were subjected to time-frequency analysis to examine stimulus-induced oscillatory responses. The epochs were imported into Matlab^®^ using the free toolbox EEGLAB (v. 9.0.5.6b) and re-referenced to the average reference. Using a custom-made scripts collection named ‘WTools” (see Parise, Csibra, and Becchio, 2013, for reference) complex Mortlet wavelets were computed for the frequencies 10-65Hz with 1Hz resolution. Total-induced oscillations were calculated, performing a continuous wavelet transformation of all epochs by means of convolution with each wavelet and taking the absolute amplitude value (i.e., the amplitude, not the power) of the results (Csibra et al., 2000). Transformed epochs were then averaged for each condition separately. To remove the distortion introduced by the convolution, we edited out 200ms at the edges of the epochs, resulting in 1200ms long segments, including 200ms before and 1000ms after stimulus onset. We used the average amplitude of the 200ms pre-stimulus window as baseline, subtracting it from the whole epoch at each frequency (Parise et al., 2013).

### Questionnaires

#### Schizotypy

The Oxford-Inventory of Feelings and Experiences-Short Form (sO-LIFE; Mason et al., 2005) assessed schizotypy dimensionality and divided the participant cohort into iSZTm/iCONm and SZTm/CONm. The sO-LIFE was chosen as the present measure of schizotypy dimensionality due to its fully dimensional approach, proposing that symptoms occurring in the schizophrenia-spectrum also occur in the typical population. The reliability of the sO-LIFE, estimated with a Cronbachs alpha of 0.79 was observed in the present cohort; illustrating a similar reliability estimate to prior literature (for example, 0.78; Fonseca-Pedrero et al., 2014). According to Mason, Linney, and Claridge (2005, p:295), the sO-LIFE has four scales: The Unusual Experiences scale contains 12 items, describing perceptual aberrations, magical thinking, and hallucinations. It is phenomenologically related to the positive symptoms of psychosis, and measures a trait often termed ‘positive schizotypy’. The Cognitive Disorganisation scale contains 11 items that parallel aspects of poor attention and concentration, as well as poor decision-making and social anxiety. It can be seen to reflect thought disorders and other disorganised aspects of psychosis. The Introvertive Anhedonia scale contains 10 items that describe a lack of enjoyment from social and physical sources of pleasure, as well as avoidance of intimacy. It can be seen to reflect weakened forms of ‘negative symptoms’, so-called negative schizotypy, or alternatively the schizoid temperament. The impulsive non-conformity scale contains 10 items describing impulsive, anti-social, and eccentric forms of behaviour, often suggesting a lack of self-control.

#### Additional Demographic Variables

A general assessment questionnaire was used to gain an overall assessment of smoking habits, hearing deficits, birth complications, and whether they, or their family have experienced mental illness. See Table 1 for the outcome of this questionnaire.

### Rationale for Analysis

The primary aim of the current research was to observe whether mothers who experience schizotypic traits demonstrate reduced neuro-oscillatory responses during a sensory gating paradigm (*Experiment 1*). As stated previously paired-tone suppression appears deficient in the schizophrenia-spectrum in the form of a smaller evoked response to *S1*, rather than larger, less effectively “suppressed” responses to *S2*. We therefore hypothesised that the high-schizotypy maternal group would illustrate reduced oscillatory amplitudes in the gamma and beta frequency ranges in response to *S1*. When exploring the dimensional nature of sensory gating abnormalities, we predicted those individuals exhibiting high schizotypy total scores would demonstrate poorer sensory gating than those who experience low schizotypy total scores. It was also hypothesised that poor sensory gating would be associated with higher manifestations of unusual experiences and cognitive disorganisation. From a theoretical perspective, poorer sensory gating suggests an inability to inhibit unimportant information, which may play a role in the poor attention experienced by these individuals. Additionally, high scores in the Unusual Experiences dimension, which describes perceptual aberrations, magical thinking, and hallucinations, which all could be explained by a lack of inhibitory control, which can be observed in poor sensory gating abilities. In addition, links have been observed between disinhibited, impulsive behaviour reflected by the impulsive non-conformity dimension and reduced sensory gating (Houston & Stanford, 2001; Liffijt et al., 2012; Park et al., 2012).

The present research also aimed to explore whether these oscillatory sensory gating responses have similar manifestations in the 6-month-old offspring of the maternal cohort (*Experiment 2*). This question was asked to explore individual differences in sensory gating abilities, in this case whether abnormalities are observable at 6-months, or whether differences are manifested later in development. Current infant literature investigating the influence of schizotypy on sensory gating development is sparse; preventing the extent to which we can state that the early manifestation of schizotypic traits could be considered as a precursor to mental illness. For this reason, it is important we explore infant electrophysiological changes that have the potential to be associated with later onset of traits.

As outlined in the introduction, responses to *S1* in the beta frequency range have been negatively correlated with the P50 suppression response to *S2* in the paired-tone paradigm (Hong et al., 2008). For this reason, the statistical analysis will predominantly focus on oscillatory responses to *S1*. Firstly, a 2 (Group: SZTm or CONm) x 2 (Paired-tone: *S1* or *S2*) x 2 (Region of Interest: right-frontal or left-frontal) repeated-measures ANOVA will be carried out to explore the maternal beta oscillatory amplitude response to *S1* and *S2*. If a significant group difference is observed then we will explore group differences specifically in *S1*. If a group by region of interest interaction is observed then we will explore whether these effects are driven by the right- or left-frontal regions. A similar strategy will be employed with the infant data.

Due to the dimensional nature of the schizophrenia-spectrum, in preference to a binary, between groups analysis, a more robust statistical approach explores the correlational relationships. Therefore, Pearson correlations will be used to explore the dimensional nature of sensory gating abilities within the range of sO-LIFE scores, including a total sO-LIFE score (total of all four dimensions), and individual dimension scores. For this element of the analysis it was hypothesised that poor sensory gating will be associated with higher levels of schizotypy, and high levels of cognitive disorganisation specifically. For the present research, a result that yields a *p*-value of less than 0.05 (*p*<0.05) have been labelled as significant.

## Results

### Experiment 1: Maternal Cohort

A 2 (group: SZTm or CONm) x 2 (Paired-Tone: *S1* or *S2*) x 2 (Region of Interest: Right-frontal or Left-frontal) x2 (Frequency: Beta or Gamma) repeated-measures ANOVA with Bonferroni Post-Hoc tests for multiple comparisons was employed. Significant main effects of paired-tone (F(1,26)=38.9, p<.001, η_p_^2^ = .60) and frequency (F(1,26)=54, p<.001, η_p_^2^ = .68) were observed. Upon further exploration of the differences in beta and gamma responses, it was observed that no significant differences were observed in gamma activity. For that reason, all analyses reported below are exploring beta activity.

A 2 (group: SZTm or CONm) x 2 (Paired-Tone: *S1* or *S2*) x 2 (Region of Interest: Right-frontal or Left-frontal) repeated-measures ANOVA explored the maternal beta oscillatory responses to *S1* and *S2*. This illustrated a significant main effect of the paired-tone (F(1,26)=31.35, p<.001, η_p_^2^ = .55) and between-subjects group (F(1,26)=8.21, p=.007, η_p_^2^ = .21). The region of interest by group interaction was also found to be significant (F(1,26)=6.57, p=.02, η_p_^2^ = .18). No further comparisons were found to be significant. To explore these group differences further, a 2 (Paired-tone: *S1* or *S2*) x2 (Group: SZTm or CONm) repeated-measures ANOVA corrected for multiple comparisons was carried out and illustrated a significant difference between the oscillatory activity towards the paired-tones (F(1,26)=18.38, p<.001, η_p_^2^ = .37), and a significant group effect (F(1,26)=16.11, p<.001, η_p_^2^ = .34). When exploring the group by paired-tone pairwise comparisons, both the SZTm and CONm demonstrate significant differences between their activity towards *S1* (*M*=.18, *SD*=.03) and *S2* (*M*=.07, *SD*=.03), but overall, the SZTm illustrated less beta activity in response to S1 (*M*=.09, *SD*=.02) and S2 (*M*=−.02, *SD*=.02), replicating previous literature (Kisley and Cornwell, 2006; Hong et al., 2008). To explore these differences further, a paired-samples t-test illustrated a significant result in the right region (*t*(27)=5.81, p<.001, 95% CI [.085, .18]); suggesting more negativity in *S2* as observed in the mother cohort (Figure 3).

**Figure 3.**
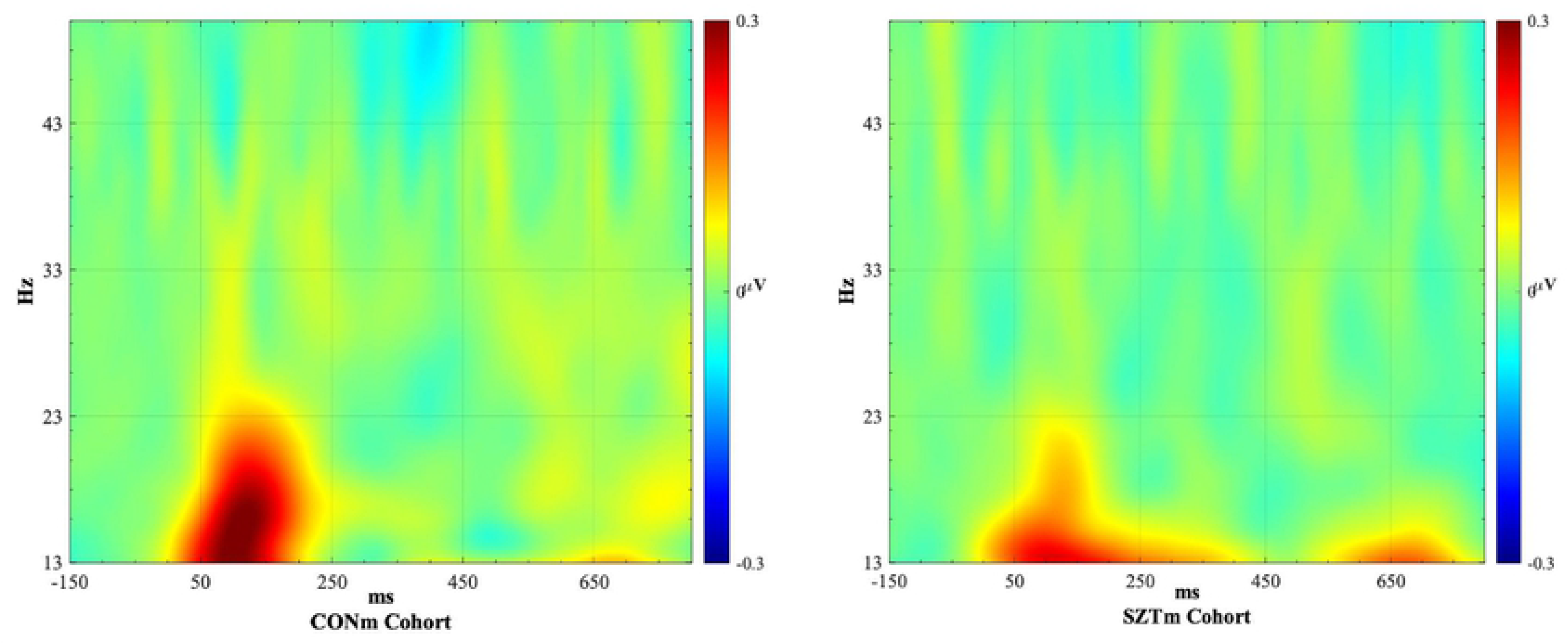
Topographical Plot of the *S1* and *S2* stimuli responses across the entire maternal cohort. Note how the amplitude related to *S1* is greater in comparison to that of *S2*.

To explore the groups beta oscillatory responses to *S1*, a one-way ANOVA was employed and demonstrated a significant between-groups effect in the right region (F(1,27)=4.18, p=.05). When exploring the descriptive statistics in the right-frontal region, the CONm illustrate a greater response to *S1* (*M*=.19, *SD*=.11) in comparison to SZTm (*M*=.11, *SD*=.09). This supports the hypothesis that mothers with high-schizotypic traits show decreased activity in response to *S1* compared to control mothers between 13-20Hz (beta; Figure 4). The present research hypothesised that the mothers with schizotypy would illustrate reduced oscillatory amplitude in response to *S1;* this hypothesis was supported by the data reported here.

**Figure 4.**
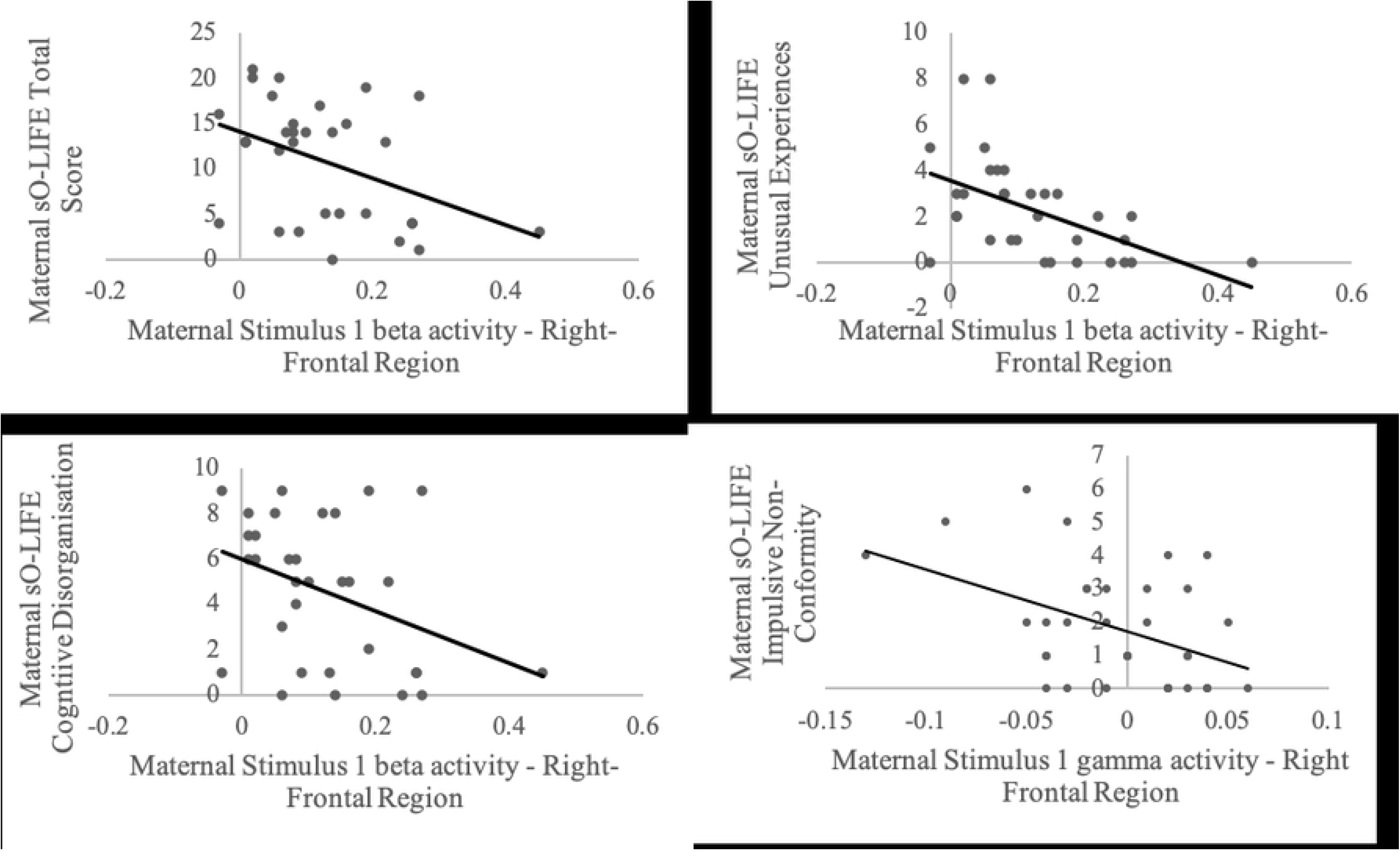
Topographical Plot of the *S1* stimuli of SZTm and CONm in the right region between 13-20Hz. Note how the CONm demonstrate greater oscillatory amplitude when compared to their SZTm counterparts whom demonstrate a similar pattern of activation but to a lesser degree.

Pearson correlations between the maternal sO-LIFE total and dimension scores and the maternal oscillatory amplitude exhibited in response to *S1* were carried out. All thirty-four maternal participants were included in this analysis. Significant negative relationships were observed between the beta responses to *S1* in the right region of interest and the sO-LIFE total (*n*=34, *r*=−.38, *p*=.03; Figure 5), and the sO-LIFE Unusual Experiences dimension (*n*=34, *r*=−.480, *p*<.005; Figure 5). Also, a strong trend towards significance was observed between the sO-LIFE Cognitive Disorganisation dimension and S1 in the right ROI (*n*=34, *r*=−.32, p=.06; Figure 5). Additionally, a significant correlation was observed between the gamma activity exhibited in response to *S1* in the right-frontal region and the Impulsive Non-conformity dimension (*n*=34, *r*=−.44, *p*<.01). These relationships suggest that a greater sO-LIFE score, which is indicative of experiencing the traits of schizotypy, the lower the oscillatory amplitude exhibited towards *S1*, supporting prior literature. No significant correlations were observed for the Introvertive Anhedonia dimension and no significant correlations were observed in the left ROI.

**Figure 5.**
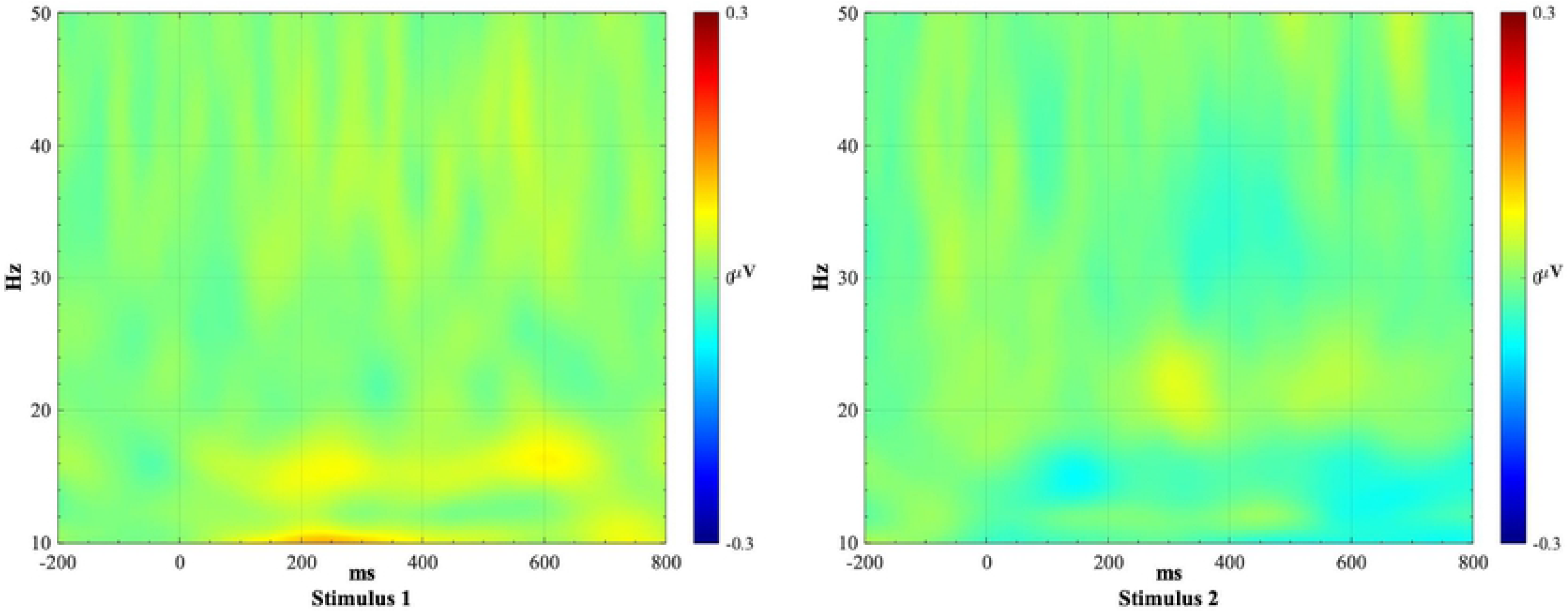
Pearson correlational analyses between the maternal sO-LIFE total and dimension scores and the maternal oscillatory activity exhibited in response to *S1* in the right-frontal regions.

### Experiment 2: Infant Cohort

#### Results

A 3 (Region of Interest: left-frontal, left-temporal, right-temporal) x 2 (Paired-tone: *S1* or *S2*) x 2 (Group: iSZTm or iCONm) repeated-measures ANOVA, with Bonferroni Post-Hoc tests for multiple comparisons, illustrated a significant difference between beta activity towards *S1* and *S2* (F(1,36)=6.89, p<.01, η_p_^2^ = .16), and a trend towards a paired-tone by region of interest interaction (F(2,72)=2.85, p=.06, η_p_^2^ = .07). No further comparisons were found to be significant.

To further explore the differences between activity in the different regions of interest, a paired-samples t-test suggested a significant difference between oscillatory amplitudes toward *S1* and *S2* (t(37)=2.82, p=.008). When exploring the group by paired-tone interaction for pairwise comparisons, it became apparent that the iSZTm did not display a significant difference between *S1* or *S2* (p=.09), but the iCONm did show a significant difference between the two stimuli (p=.04); this does not, however, drive a significant group effect. A paired-samples t-test was undertaken to illustrate a significant difference between the frequencies observed in the right-temporal region (t(37)=2.82, p=.008, 95% CI [.02,.13]); illustrating decreased activity in *S2* compared to *S1* (Figure 6). No further significant effects were observed. We hypothesised that if the infants were to demonstrate a similar deficit to their biological mothers, then they too would exhibit reduced oscillatory amplitudes in the gamma and beta frequency ranges in response to *S1*. The current data, however, did not show these deficits in the infant cohort, supporting the results of Hong et al. (2008) in that typical first-degree relatives of those diagnosed with schizophrenia display no difference in oscillatory gating compared to controls.

**Figure 6.**
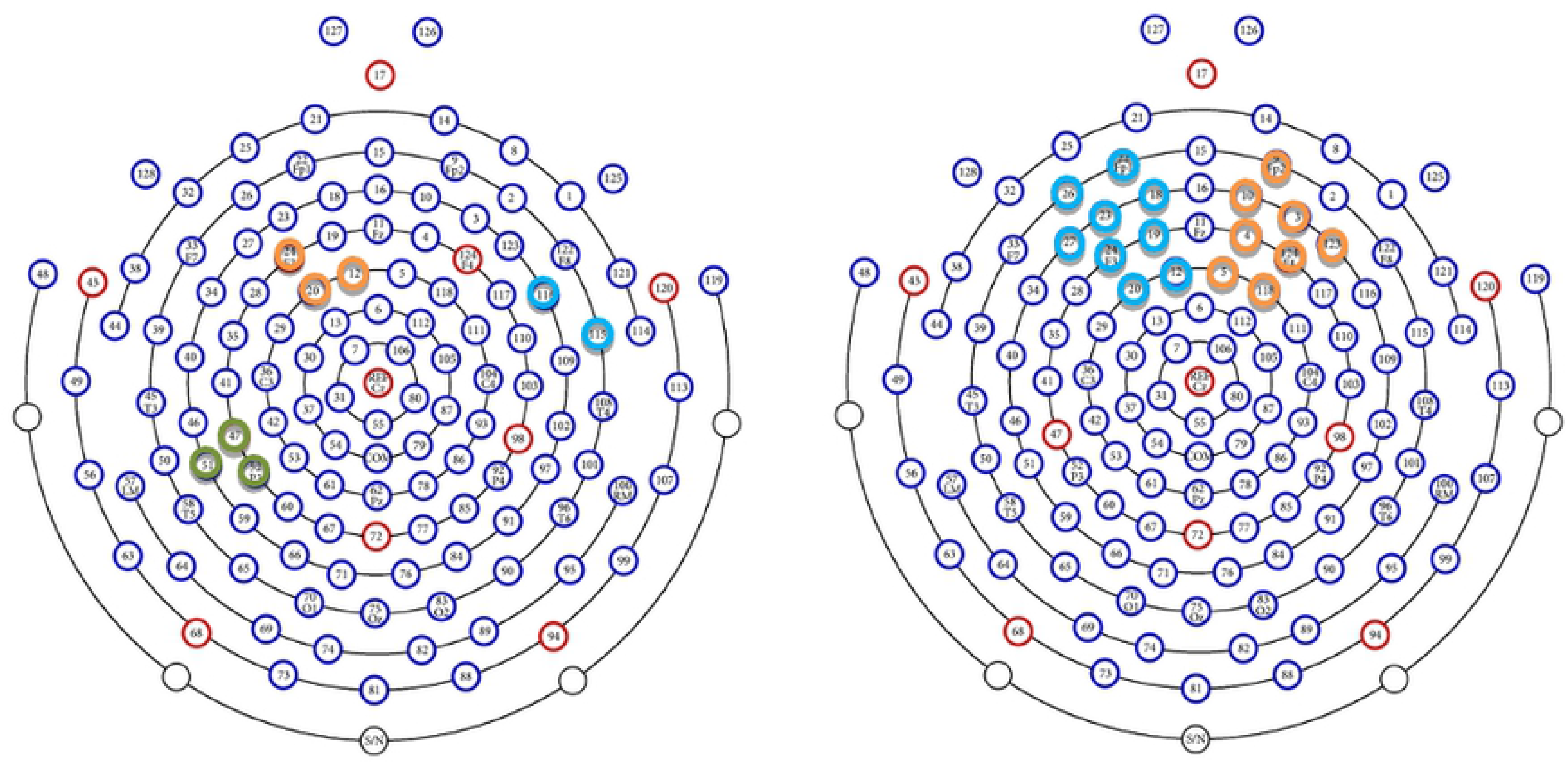
Topographical Plot of *S1* and *S2* across the entire infant cohort. Note how the amplitude related to S1 is greater in comparison to that of *S2*. A greater amplitude can be observed between 10-20Hz across the time-window of *S1*.

For all the infants who took part in *Experiment 2* (*n*=75), sO-LIFE scores were collected from their Mother, regardless of whether they were invited to participate in *Experiment 1* (*n*=34) themselves. As a result of this, the correlational analyses discussed here refers to the relationship between the infant’s oscillatory activity (*n*=46) and their own mother’s sO-LIFE score (*n*=46); therefore, exploring the relationship between the schizotypy dimensionality in the infant-mother dyad and its influence on the infant’s oscillatory activity. The research was designed in this way as it is not possible to reliably infer schizotypy dimensionality in infants, therefore their status was inferred through the infant-maternal dyad (Keshaven et al., 2008), from their mother’s sO-LIFE score.

Pearson’s correlations explored maternal sO-LIFE total and dimension scores (associated with each infant-maternal dyad) and the corresponding infant’s oscillatory activity exhibited in response to *S1* (*n*=46). No significant relationships were observed in any of the left-temporal, right-temporal, or left-frontal regions of interest.

## General Discussion

We hypothesized that mothers in the schizotypy group would exhibit reduced oscillatory activity in response to *Stimulus 1* during the sensory gating paradigm when compared to their control counterparts. The data supported this notion by demonstrating how mothers with high-schizotypy displayed reduced beta activity towards *S1*, replicating the previous literature (Johannesen et al., 2005; Brenner et al., 2009; Hall et al., 2011). When exploring this question in a more dimensional manner, it was hypothesised that those mothers exhibiting high schizotypy would display poorer sensory gating than those who experience low schizotypy scores; particularly in higher manifestations of cognitive disorganisation and impulsive non-conformity. Correlational relationships suggested that reductions in activity toward *Stimulus 1* were observed in both beta and gamma activity among those mothers who exhibited higher schizotypy scores; specifically, the sO-LIFE total score, impulsive non-conformity, cognitive disorganisation, and the unusual experiences dimension.

The present research also explored whether these oscillatory responses had similar manifestations in the 6-month-old offspring of the maternal cohort. The infant cohort showed no significant differences between the groups; suggesting that having a mother with schizotypy did not influence the infants’ oscillations in relation to sensory gating at the group and dimensional level. This finding supports that reported by Hong et al. (2008) who described how first-degree relatives of those on the schizophrenia-spectrum displayed no difference in their event-related oscillation gating responses when compared to controls.

Despite Hong et al. (2008) describing how sensory gating oscillatory responses were not shared between the schizophrenia-spectrum and first-degree relatives, contradicting evidence suggests that neuro-oscillatory deficits are observed in the first-degree relatives of schizophrenia patients, who exhibit schizotypic-personality traits (Hong et al., 2004). The present research, however, supports the idea that, at least at 6-months of age, oscillatory deficits are not influenced by their mother experiencing schizotypic traits. Existing infant literature investigating the influence of parental schizotypy on sensory gating development is sparse; thus all predictions made were based on other measures of maternal disposition. For example, infants of mothers reporting less ‘social recognition’ and higher ‘social isolation’ developed less pre-pulse inhibition during the four months after birth than did infants of mothers with higher ‘social recognition’ and less ‘social isolation’ (Huggenberger et al., 2013). This confirms that parental disposition can impact the inhibitory abilities of their infants at 4-months. With the knowledge it is not that parental disposition can influence infants’ inhibitory abilities at 4-months, it is of fundamental interest to explore the different electrophysiological frequencies associated with auditory sensory gating to explore whether these endophenotypic deficits are observed earlier in the developmental trajectory. Perhaps inhibitory abilities may be influenced by parental schizotypy later in development, but exposure at 6-months postnatal does not demonstrate individual differences.

It should be articulated that schizotypy, for the purpose of the current research, was defined using the sO-LIFE, with mothers classed as schizotypic if their sO-LIFE score was half a standard deviation above the mean; weighing in favour of a fully-dimensional approach and replicating the approach taken by Park et al. (2015). However, this approach could limit our understanding of schizotypy as a construct with separable and well-identified components (Kwapil, Barrantes-Vidal, and Silvia, 2008). To account for this, the present research included correlational analyses between the sO-LIFE total and dimension scores in comparison to the oscillatory amplitude toward *S1*. This found that mothers with greater sO-LIFE scores: an indicator of schizotypy dimensionality, illustrated lower oscillatory activity towards *S1*. The literature differs in terms of which dimensions of schizotypy are associated with deficits in sensory gating. For example, it has been reported to be the negative dimension (introvertive anhedonia; Wang et al., 2004), the positive dimension (unusual experiences; Croft et al., 2001, 2004), and the cognitive disorganisation dimension (Evans et al., 2007; Park et al. 2015). The literature therefore provides support for associations with multiple dimensions of schizotypy. The present research therefore illustrated a strong trend towards a negative relationship between poor sensory gating and the cognitive disorganisation dimension, a negative relationship between poor sensory gating and unusual experiences and the impulsive non-conformity dimension. This is in support of research by Park et al. (2015) who found attenuated P50 suppression was observed among a group with high impulsive non-conformity scores as a link has been made between disinhibited, impulsive behaviour and reduced sensory gating (Houston and Stanford, 2001; Liffijt et al., 2012). These correlational relationships were not however, observed in their infants in the present research.

A strength of the present research is in the multiple electrode sites utilised for analysis when contrasted to prior research, which used low-density montages or singular electrodes (for example, CZ, Hong et al., 2004). The present research also highlights the complexity of recording electrophysiological activity during sleep where infants produce unpredictable movements; increasing artefacts and attrition rates for inclusion in the final analyses. A limitation of the present research is the use of all white-Caucasian sample, which limits the generalisability of the results. Thus, future exploration of similar concepts should aim to employ a more varied cohort. A further limitation of the present study is the lack of previous literature observing sensory gating time-frequency oscillations with infants. This is the first time this has been demonstrated, to our knowledge. For this reason, there is little literature on which to base our predictions and have a priori topographical hypotheses; all topographical predictions and outcomes must therefore be interpreted with caution. Future analyses should explore first-degree relatives during childhood to explore at what timepoint these schizotypic deficits develop.

Decomposing specific frequency components associated with event-related potentials is a key step in understanding sensory gating as each event-related oscillation is thought to reflect different aspects of early sensory information processing (Kopell et al., 2000; Traub et al., 1999; Traub et al., 1999; Basar-Eroglu et al., 1996). For example, beta activity has been implicated in novelty detection and salience encoding (Uhlhaas et al., 2008; Kopell et al., 2000; Kisley and Cornwell, 2006), whereas gamma oscillations are associated with the immediate registration of sensory stimuli (Kopell et al., 2000; Tallon-Baudry and Bertrand, 1999) reflecting the “binding of spatially distinct neuronal activities within and between brain regions” necessary for coherent sensory perception and processing (Uhlhaas et al., 2008; Singer and Gray, 1995). Hall et al. (2011) proposed that the components of information processing assessed by gamma and beta gating appear to be independent from those mediated by P50 suppression. This stems from the P50 gating deficit found in first-degree relatives in contrast to their typical gamma and beta gating responses. They concluded that this combination of ERP and oscillatory findings suggests that P50 sensory gating is functionally distinct from frequency-specific gating responses. This conclusion is consistent with the results of Clementz and Blumenfeld (2001) who illustrated a significant P50 gating deficit in schizophrenia patients but typical gamma gating, and the independence of a significant theta/alpha-band gating deficit and P50 gating in schizophrenia patients and their relatives, which was observed by Hong et al (2008). Moreover, Smith et al. (2019) illustrated correlational relationships between 6-month-old infants’ P50 gating responses and their mothers sO-LIFE score, suggesting that infants of schizotypic mothers were more likely to display reduced suppression compared to controls. However, the present research does not show the same deficit in first-degree relatives.

The present research supports the notion that those individuals who display schizotypic traits display differences in oscillatory function, alike to those displayed by patients with schizophrenia, when contrasted with controls, however, the infant first-degree relatives of these individuals do not present with the same deficits. Psychiatric disorders, such as schizophrenia, may therefore be thought of as a ‘shift’ in the continuous distribution of neurodevelopmental traits towards greater impairment, whilst maintaining overlap with the population distribution (Hengartner and Lehmann, 2017). Establishing brain-behaviour links in both clinically significant behaviours and those of typical development is an important step in further understanding the continuities and discontinuities that exist between typical and pathological behaviour.

The present research has illustrated how mothers with sub-clinical schizotypy who are part of the general population display reduced beta activity towards *Stimulus 1* in a paired-tone paradigm; which parallels previous literature exploring this same mechanism in other groups located on the schizophrenia-spectrum. This shared deficit supports the potential contribution of oscillatory abnormalities of this kind as an endophenotype for the schizophrenia-spectrum, although further exploration and replication is needed. In contrast to the maternal cohort, the infants demonstrated no group differences, highlighting that having a mother with schizotypy did not influence their oscillatory sensory gating abilities at 6-months-old. The establishment of this method of measuring sensory gating in infants (Kisley et al., 2003) is crucial as it increases our understanding of the developmental timeline of these disorders, will perhaps allow for the development of novel prevention strategies, and assist in the exploration of endophenotypic deficits observed earlier in the developmental trajectory.

## Author Contributions

ES: Conceptualisation, Methodology, Investigation, Formal Analysis, Writing - Original Draft, Writing - Review & Editing. TJ: Supervision, Writing - Review & Editing. VM: Supervision, Writing - Review & Editing.

## Acknowledgements

This work is supported by the Leverhulme Trust Doctoral Scholarship Programme in Interdisciplinary Research on Infant Development. The authors wish to acknowledge Dr Eugenio Parise who assisted in the analysis of the data through the use of his MatLab custom-made scripts collection named ‘WTools’. The authors also wish to acknowledge the involvement and support of the families who participated in the present research.

